# Proteomics, lipidomics, metabolomics and 16S DNA sequencing of dental plaque from patients with diabetes and periodontal disease

**DOI:** 10.1101/2020.02.25.963967

**Authors:** Katherine A. Overmyer, Timothy W. Rhoads, Anna E Merrill, Zhan Ye, Michael S. Westphall, Amit Acharya, Sanjay K. Shukla, Joshua J. Coon

## Abstract

Oral microbiome influences human health, specifically pre- and type 2 diabetes (Pre-DM/DM) and periodontal diseases (PD), through complex microbial interactions. To explore these relations, we performed 16S rDNA sequencing, metabolomics, lipidomics, and proteomics analyses on supragingival dental plaque collected from individuals with Pre-DM/DM (n=39), Pre-DM/DM and PD (n=37), PD alone (n=11), or neither (n=10). We identified on average 2,790 operational taxonomic units and 2,025 microbial and host proteins per sample and quantified 110 metabolites and 415 lipids. Plaque samples from Pre-DM/DM patients contained higher abundance of *Fusobacterium* and *Tannerella* vs. plaques from metabolically healthy. Phosphatidylcholines, plasmenyl-phosphatidylcholines, ceramides containing non-OH fatty acids, and host proteins related to actin filament rearrangement were elevated in plaques from PD vs. non-PD. Cross-omic correlation analysis enabled the detection of a strong association between *Lautropia* and mono-methyl phophospotidlyethanolamine (PE-NMe), striking because synthesis of PE-NMe is uncommon in oral bacteria. Lipidomics analysis of *in vitro* cultures of *Lautropia mirabilis* confirmed the bacteria’s synthesis of PE-NMe. This comprehensive analysis revealed a novel microbial metabolic pathway and significant associations of host-derived proteins with PD.

## Introduction

The human oral cavity harbors a wide variety of microbes - over 700 species ^1^- and has some of the highest microbial diversity observed in humans ^2^. These oral-associated bacteria reside in saliva, on the tongue and cheeks, and in biofilms on tooth surface and under the gum lining ^2^. The development of plaque biofilms is particularly important to the etiology of oral diseases, like tooth decay and periodontal disease (PD) ^3,4^. And importantly, the pathogenic oral microbiota that contribute to the progression of PD are also correlated with systemic diseases, including diabetes, arthritis, and heart disease ^5-7^, suggesting that oral microbial ecologies have a broad impact on human health, and a better understanding of pathogenesis and host-microbe interactions will be essential for mitigating negative effects of pathogenic microbiota.

With poor oral hygiene, bacterial populations accumulate, become increasingly diverse, and cause gum inflammation ^8^. The progressive shifts in the plaque biofilm diversity are strongly associated with PD incidence and severity ^4,9^. In particular, species that form what is called the ‘red complex’, *Tannerella forsythus, Porphyomonas gingivalis*, and *Treponema denticola*, are associated with gum bleeding on probing and probe depth, two common markers of PD severity ^9^. These red complex microbial populations are observed in conjunction with species like *Fusobacterium nucleatum, Prevotella intermedia, Prevotella nigrescens, Peptostreptococcus micros* species, which are more mildly associated with PD ^9^ and often are observed in biofilms before the red complex.

During the development of dental plaque, increases in microbial diversity are mediated by changes in the microenvironment and manifestation of microbial niches ^10,11^, as the local environment becomes optimal to support population growth. This growth can be aided by microbe-microbe interactions, host-microbe interactions, and metabolite availability ^4^. As dental plaque biofilms become established, measurable changes in microbial abundances as well as metabolites and host-factors occur. Thus, a holistic approach to studying dental plaque could provide insight into how microbial populations interact and how they associate with host health.

Frequently 16S/18S rDNA sequencing is employed to estimate size and diversity of microbial populations ^1,2^; however, this method offers little information about microbial function or local environmental factors, although researchers have come up with ways to use 16S sequencing to deduce some functional information (i.e., PICRUSt) ^12^. More recently metagenomics and metatranscriptomics approaches have afforded greater evidence for microbial functional potential, *i.e.*, what genes are present and/or expressed in the population ^13^. These methods can provide clues to how microbes might interact within the biofilm, but importantly, these conclusions are greatly strengthened by biomolecule measurements, for example, when metagenomics is paired with metabolomics ^14,15^. Mass spectrometry (MS)-based ‘omic analyses – like metabolomics, lipidomics, and proteomics – uniquely offer high-throughput means of assessing the molecular details of the local environmental niches these microbes occupy as well as information about community composition and functional-level information. Indeed, several studies have used MS-based ‘omics to assess the oral microbiome, focusing on either metaproteomics ^16,17^ or metabolomics ^18-21^. As of yet, discovery lipidomics is not being applied to study the oral microbiome, and generally, the use of multi-omics for studying the oral microbiome is still uncommon ^21^, despite the fact that it has potential to offer a wealth of information about the local microbe-host environment ^22^.

We leveraged 16S rRNA gene sequencing and high-resolution MS to profile microbes, proteins, lipids, and metabolites in human dental plaques from nearly 100 individuals with PD and/or pre-diabetes/diabetes (Pre-DM/DM). Altogether we performed >650 GC or LC-MS/MS experiments, collected >4.5 million tandem mass spectra, and on average identified several thousand biomolecules in each sample. Using ecology diversity metrics, we identified increases in microbial diversity and changes in the microbial populations with PD and Pre-DM/DM, suggestive of microbial dysbiosis occurring with these diseases. Further, we identify hundreds of proteins, metabolites, and lipids associated with PD and/or Pre-DM/DM. Based on these findings, we describe a rare lipid synthesis pathway in one of the common oral bacteria (*Lautropia mirabilis*) and demonstrate that these compounds along with other microbial-molecule associations can link microbial populations to function.

## Results

We collected three supragingival plaque samples from buccal and palatal tooth surfaces from each of the 97 study participants (Pre-DM/DM, n=39; Pre-DM/DM with PD, n=37; PD, n=11; or neither, n=10, **Table 1**). The study participants were primarily non-Hispanic white (93%) and non-smokers (84%); the Pre-DM/DM participants were significantly older than the metabolically healthy participants (62±16 vs. 43±16 years old, p<0.001), **Table 1**. One plaque sample was used for 16S rDNA sequencing, and two plaque samples were used for MS-based analyses – proteomics, lipidomics, and metabolomics (**Figure 1**). For 16S sequencing, DNA sequences are easily amplified to enhance the quantitative signal, and data processing methods for determining the features present (*i.e.*, microbial populations) are well established. In contrast, for the MS-based approaches two major challenges - limited sample amount and feature identification - still persist. To address them, we maximized our limited samples by extracting several compound classes (small-molecules, lipids, and proteins) from a single plaque sample and used comprehensive libraries of standards and databases to annotate our raw data (>650 raw files). This methodology enabled detection of 50,752 operational taxonomic units (OTUs by rRNA sequencing; 99% sequence similarity; ∼ 2,790/sample), 12,346 annotated protein groups (∼ 2,025 proteins/sample), 415 lipids, and 89 metabolites (**Table S1, Figure 1**), making this study the most comprehensive analysis of dental plaque to date.

**Table 1.** Patient population statistics. Patients were grouped by pre-diabetes/diabetes (Pre- DM/DM) and periodontal disease (PD) status. Pre-DM/DM patients were significantly older and had higher HbA1c and fasting blood glucose (p<0.05).

| Group | n | Sex | Age* | Race/ethnicity |  | HbA1c* | Fasting blood glucose* | Periodontal disease | Tobacco use |
| --- | --- | --- | --- | --- | --- | --- | --- | --- | --- |
|  |  | (female/<br>male) | Years<br>Mean<br>(St.Dev.) | White | Hispanic | %<br>Mean (Std.Dev.) | mg/dL<br>Mean (St.Dev.) | (moderate/<br>severe) | (current/<br>former) |
| Pre-DM/DM + PD | 39 | (22/17) | 61.3(15.7) | 37 | 3 | 7.1(2.1) | 126.0(45.4) | (35/4) | (11/10) |
| Pre-DM/DM | 37 | (20/17) | 64.1(16.3) | 36 | 1 | 6.7(1.6) | 126.1(37.9) | (0/0) | (3/17) |
| PD | 11 | (9/2) | 45.2(15.9) | 10 | 0 | 4.7(0.2) | 91.3(6.6) | (9/2) | (2/6) |
| Healthy (non-PD/non-DM) | 10 | (8/2) | 39.6(16.0) | 10 | 0 | 4.9(0.1) | 91.2(6.1) | (0/0) | (0/4) |

\* $p < 0.05$ , Pre-DM/DM vs. non-Pre-DM/DM

**Figure 1.**
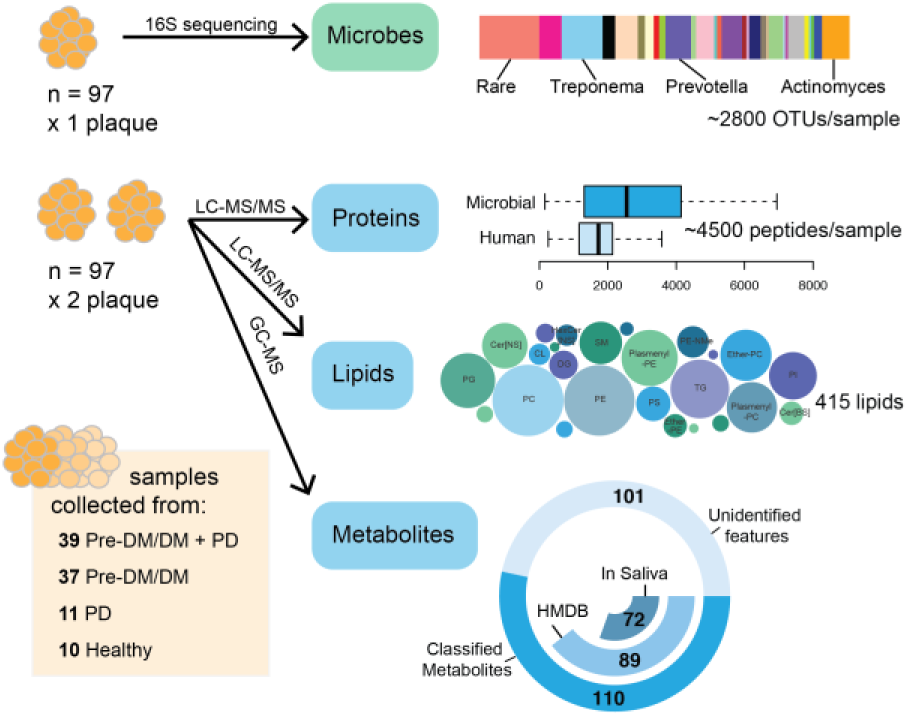
Sample collection and processing strategy for the microbiome, proteome, lipidome, and metabolome analyses. Patients were classified by pre-diabetes/diabetes (Pre- DM/DM) and periodontal disease (PD) status. One plaque sample was used for 16S rRNA sequencing to generate a list of operational taxonomic units (OTUs), and 2 plaque samples were used for mass-spectrometry based analyses - proteomics, lipidomics, and metabolomics, which led to the identification of ∼4500 peptides, 415 lipids, and 126 metabolites per sample, respectively.

To permit further informatic analysis, we assembled the dataset by first normalizing lipidomics and metabolomics data to account for sample-size variation, then applying batch-normalization ^23^. Upon quality assessment, we removed two proteomic samples due to the low number of identified features. Finally, as we structured our statistical analysis to compare the plaque composition in healthy vs. PD or Pre-DM/DM samples, we accounted for confounding variables, such as age, sex, and tobacco use. In addition to deciphering features associated with the disease, we assessed microbiota diversity using both 16S rRNA-based taxonomy and proteomics-based taxonomy, determined molecular composition of the plaque biofilms, and clustered co-occurring molecules to infer molecular associations.

### Microbial populations show unique dysbiosis with PD and Pre-DM/DM

Across our samples we observed hundreds of common OTUs (256, >75% of patients) and numerous rare OTUs (50,496, <10% of patients). We used these to calculate standard ecology diversity metrics. The diversity within a sample, known as alpha diversity, varied across the plaque samples (**Figure 2a**) and was greater with disease states (log-likelihood ratio test PD vs. non-PD, p = 0.03, and Pre-DM/DM vs. metabolically healthy, p = 0.004). Similarly, we assessed the diversity across samples or beta diversity using the Bray-Curtis distance (**Figure 2b**). In contrast to alpha diversity, beta diversity was less associated with the diseases, as we observed a small but significant effect with Pre-DM/DM on beta diversity (permutation MANOVA, R^2^ = 0.017, p = 0.03). These data are consistent with previous publications reporting that biodiversity tends to be elevated in oral disease ^24-26^.

**Figure 2.**
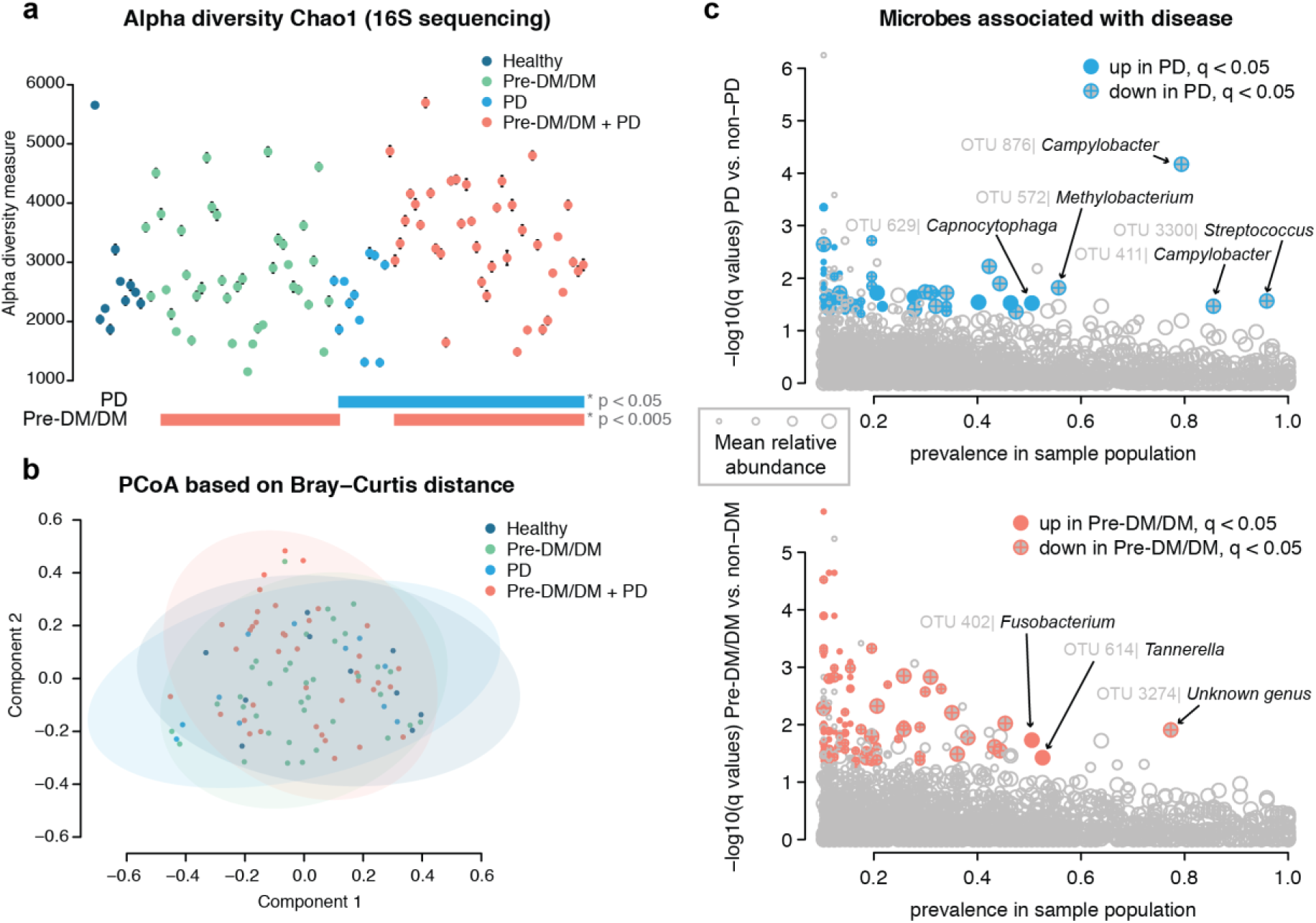
Diversity of microbial populations is similar across patient plaque samples. Patients’ plaque microbial communities were assessed by 16S rRNA sequencing. The Chao1 Index was varied across patients (a) and were significantly different between patients with periodontal disease (PD) vs. non-periodontal disease and pre-diabetics/diabetics (Pre- DM/DM) vs. non-diabetics (log-likelihood ratio test, p = 0.03 and p = 0.004, respectively). The Bray-Curtis distance for measuring beta diversity showed no significant difference between groups (b). When specific operational taxonomic units (OTUs) were assessed for association with PD (c, above) or Pre-DM/DM (c, below), we found several OTUs that were detected in a majority of our samples (prevalence in sample population > 50%) that also had q-values of < 0.05 and log2 fold-change greater than 1 (up) or less than −1 (down) in disease vs. non-disease.

To establish which microbial species were significantly associated with disease, we performed linear regression analysis on OTUs observed in >10% of the samples (n = 4169, 3048 with genera assignment, 29 with genera and species assignment). We filtered our results based on disease significance, log_2_ fold-change (Log_2_FC), and prevalence in the sample population (**Figure 2c, Table S1**). This data filtering led us to identification of 56 microbial species that were significantly different between PD and non-PD samples and 36 that were significantly different between Pre-DM/DM and metabolically healthy conditions. In PD we observed a lower relative abundance of several *Streptococcus spp., Campylobacter spp., Actinomyces spp.*, and a *Methylobacterium sp.* Lower levels of *Streptococcus spp.* in PD have been reported previously ^27,28^, and the loss of this genera is believed to play a role in disease progression by creating space for more pathogenic bacteria to thrive ^27^. We also observed higher abundance of *Capnocytophaga* in plaques of PD patients (**Figure 2c, Table S1**).

Comparing Pre-DM/DM vs. metabolically healthy group, we identified 36 significantly different OTUs (**Figure 2d, Table S1**). Specifically, microbes belonging to *Fusobacterium* and *Tannerella* genera - classic periodontal pathogens ^9^ - were elevated in Pre-DM/MD. In agreement with our findings, a recent study reported that red complex genera, which includes *Tannerella*, were especially elevated on healthy tooth sites in DM vs. non-DM ^29^. The elevation of these pathogenic bacteria in supragingival plaque could indicate an overall dysbiosis in DM, potentially increasing propensity for PD. Overall, our data support the hypothesis that dysbiosis occurs in both PD and Pre-DM/DM and importantly, that the microbial changes that take place in either disease state are distinct.

### Metabolites, lipids, and proteins found in plaque are of human and microbial origins

To further characterize the composition of the oral plaques, we looked to our MS-based ‘omics data. We measured thousands of compounds, including amino acids, monosaccharides, phospholipids, triglycerides, and human and microbial proteins (**Figure 1**). For each of the MS-based ‘omics intra-patient abundance measurements were more similar than inter-patient ones (**Figure S2**). As expected, the identified compounds were likely of both human and microbial origins. A majority of the metabolites identified by GC-MS had been previously identified in saliva (88%, 72 of 89 compounds), as annotated in the Human Metabolomics Database (HMDB) ^30,31^. The remaining 17 compounds had been detected in human feces or were annotated as human cellular metabolites, indicating a probable human, microbial, or food origin.

The lipids consisted of phospholipids – primarily phosphatidylcholine (PC), phosphatidylethanolamine (PE), and phosphatidylglycerol (PG) – as well as triglycerides and ceramides. Approximately 28% of the lipids contained odd-chain fatty acyl tails (**Figure S3**), which are more commonly found in bacteria than eukaryotic cells ^32^. The percentage of odd-chain acyl tails was higher in the PGs (∼45%) than in other lipids, likely because PGs are also more common in bacterial than in mammalian membranes ^33^.

Finally, the proteins identified in the plaques belonged to various taxonomic branches. Nine percent were from eukaryotic taxonomic branches, and likely human in origin, and 15% were unassigned or assigned to the root taxonomy level (**Table S1**). A majority of the proteins were from various bacterial genera, including *Actinomyces, Corynebacterium, Leptotrichia, Capnocytophaga*, and *Prevotella* (top-5 genera based on representative proteins). *Actinomyces* and *Corynebacterium* were previously found to contribute to a majority of the oral biofilm proteome ^34^, suggesting that these genera indeed make up a majority of plaque proteins.

### Proteomics and 16S data provide complementary information about microbial diversity

Similar to 16S data, proteomics data provided insight into microbial diversity and relative abundance of genera. According to proteomics measurements, alpha diversity tended to be higher in plaques from PD and Pre-DM/DM as compared to heathy patients - a similar finding to what was observed with 16S data. After accounting for confounders (sex, age, and smoking status) this effect was, however, reduced (**Figure S4**). Beta diversity showed no significant effect with PD or Pre-DM/DM. Overall, the proteomics data resulted in similar trends, but with lesser effect sizes, to those observed in the 16S data.

Next, we compared how well OTU-based bacterial identification matched to the bacterial identification determined by proteomics. We calculated genera and phyla overlap (49 and 10, respectively, **Figure 3a-b**) and compared average relative abundance of genera between the two methodologies (**Figure 3c-d**). Note that different taxonomic ontologies were used for 16S data (Greengenes ^35^) and proteomics (NCBI taxonomy ^36^), and we, therefore, expected differences in taxonomy assignments ^37^. Despite this, we observed considerable overlap (∼34%) between all detected genera and even greater similarity for common genera (∼89% overlap between genera detected in >90% samples). Additionally, abundance measurements at the genus level exhibited good agreement, as the observed within-individual correlation of genera abundance between 16S data and proteomics was better than expected by chance alone (**Figure 3c**, p < 0.001). We directly compared the most abundant genera detected by the two methods across our samples (mean relative abundance > 0.1%, **Figure 3d**); the notable differences between the methods were greater relative abundance of *Actinomyces* and *Corynebacterium* in proteomics vs. 16S data and greater relative abundance of *Prevotella, Selenomonas, Streptococcus*, and *Veillonella* in 16S vs. proteomics data.

**Figure 3.**
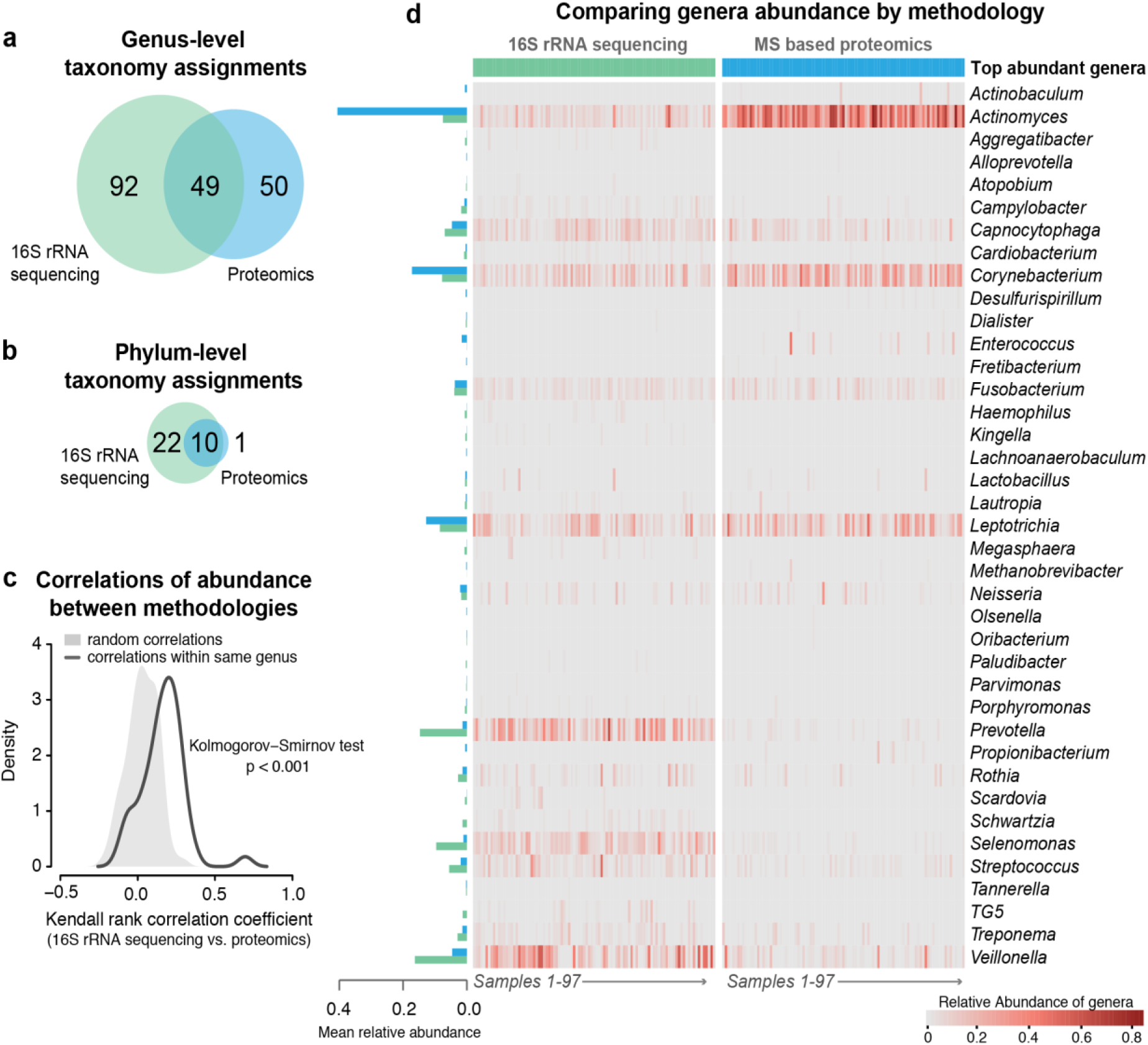
Proteomics and 16S sequencing approaches result in similar taxonomic assignment across patient samples. Taxonomy assignment resulted in 141 genera and 31 phyla by 16S rDNA sequencing and 99 genera and 11 phyla by proteomics, with 49 genera and 10 phyla in common (a and b). Correlation between genera abundance by 16S rDNA sequencing vs. proteomics within individuals was better than expected by chance alone (c). The top-abundance genera showed good overlap (d), except *Actinomyces* and *Corynebacterium* were found in greater abundance by proteomics, while *Prevotella, Selenomonas, Streptococcus*, and *Veillonella* were found in greater abundance by 16S rDNA sequencing.

### Plaques from PD patients contained elevated human-derived proteins, PCs, and amino acids

We found 234 proteins, 76 lipids, and six metabolites that exhibited significant associations with PD, and 46 proteins, six lipids, and one metabolite that significantly associated with Pre-DM/DM (q-values < 0.05, **Table S1, Figure 4**).

**Figure 4.**
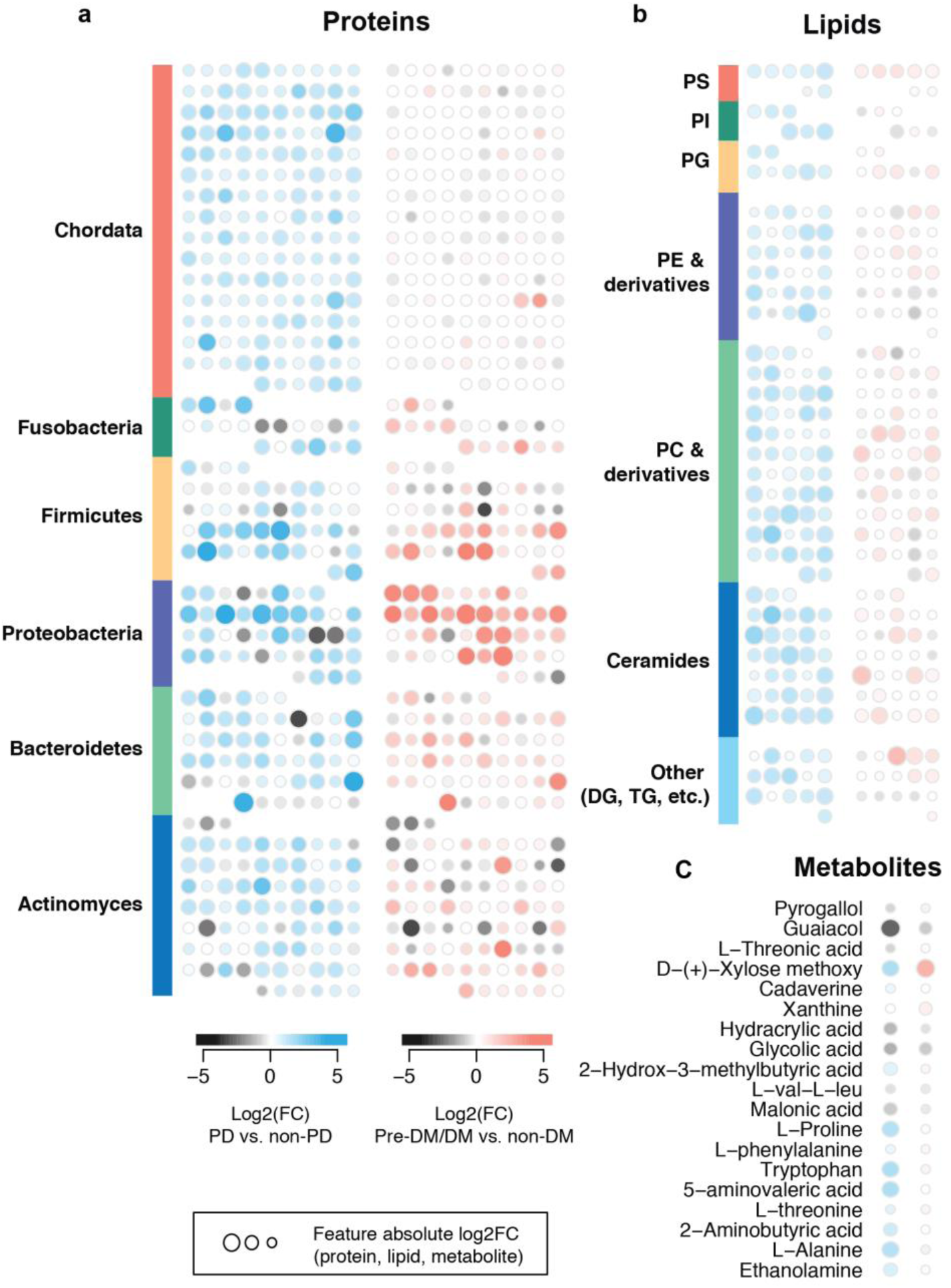
Comparison of biomolecule abundance changes occurring with periodontal disease (PD) and pre-diabetes/diabetes (Pre-DM/DM). Each symbol represents a unique protein (a), lipid (b) or metabolite (c). Proteins are grouped by phylum (a) and lipids are grouped by lipid class (b). PS, phophospotidlyserine; PI, phophospotidlyinositol; PG, phosphotidlyglyercol; PE, phospotidlyethanolamine; PC, phosphotidylcholine; DG Diacylglycerides; TG, triglycerides; PD, periodontal disease; DM, diabetes.

For the plaque proteins that were associated with either PD or Pre-DM/DM (234 and 46, respectively), we assessed protein enrichment for certain taxonomic classes or functional pathways (KEGG, GO). Proteins that were elevated in plaque from patients with PD were enriched in proteins from genera *Oribacterium* (q-value < 0.05) and *Homo* (q-value < 0.01). PD-associated proteins were also more enriched for GO terms for signaling, actin cytoskeleton organization, cell communication, and NF-kappaB transcription factor activity (q-values < 0.002). Proteins that were elevated in plaques from Pre-DM/DM were enriched in proteins from *Campylobacter* (q-value < 0.02). Overall, PD coincided with greater detection of human-derived proteins (phylum Chordata) than Pre-DM/DM (**Figure 4a**) and suggests potential host-factors relevance in PD. Consistently, prior metaproteomics studies found human saliva proteins related to innate immunity to be elevated in periodontitis ^17^.

Many lipids were elevated in plaques from PD vs. non-PD (n=76), and this list was over-represented in PCs (n=15), plasmenyl-PCs (n=11), and ceramides containing non-OH fatty acids and sphingosines (Cer[NS], n = 11). Some of these lipids include Cer[NS] d18:0_15:0, Cer[NS] d18:1_15:0, PC 39:2, and plasmenyl-PC P-18:0_20:4. Comparing lipids in Pre-DM/DM vs. metabolically healthy samples, we detected five lipids elevated with Pre-DM/DM (TG 42:1, Cer[NS] d16:0_15:0, ether-PC O-32:1, ether-PC O-40:4, and TG 12:0_12:0_12:0) and one lipid that was lower in Pre-DM/DM (PC 18:2_18:1). Overall, PD resulted in larger changes in plaque PCs and ceramides than Pre-DM/DM (**Figure 4b**).

Metabolites 5-aminovaleric acid, L-alanine, tryptophan, L-proline, and D-xylose were elevated in PD vs. non-PD. D-xylose was also elevated in Pre-DM/DM. One metabolite, glycolic acid, was reduced in PD-associated plaques. Comparison of these metabolite changes in PD vs. Pre-DM/DM is shown in **Figure 4c**; overall, PD resulted in elevated plaque amino acids, which is consistent with prior studies showing elevated amino acids in saliva of PD patients ^20,38,39^.

### Associations between lipids, metabolites, and proteins indicate disease signatures related to actin-filament rearrangement

To explore how the detected molecules might relate to the microbial populations, we performed correlation analysis across datasets using the Kendall non-parametric test. We found hundreds of significant correlations and strikingly, distinct clusters of correlations between proteins and metabolites and lipids, indicating that groups of proteins were related to specific metabolite/lipid profiles (**Figure 5a**). We used hierarchical clustering with k-means k=8 to define protein clusters and k=6 to define metabolite/lipid clusters. Several protein clusters (clusters 2 and 3) contained a large portion of the disease-associated proteins (**Figure 5b**).

**Figure 5.**
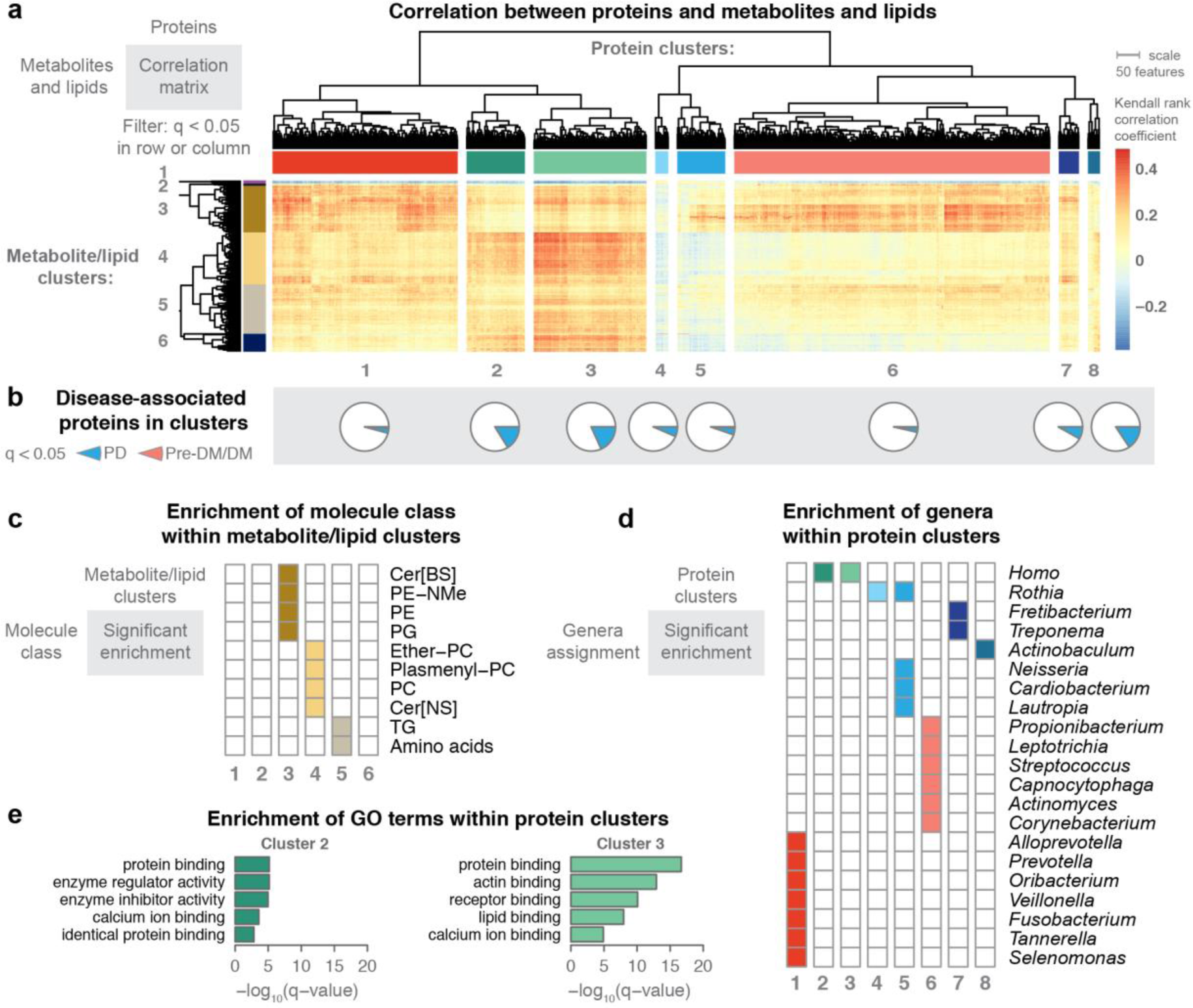
Metabolite and lipid associations with plaque proteins manifested genera- specific clusters. MS-acquired data from two plaque samples per patient were used to investigate how metabolite and lipid signatures correlate with plaque proteins. Kendall rank-based correlation was used to filter associations; metabolites, lipid, or proteins with at least one significant association (q < 0.05) are presented in the heatmap with hierarchical clustering of rows and columns (a). Using k-means (k=8) to define protein clusters, we observed disease associated-proteins in each cluster (b), but clusters 2 and 3 had higher proportions of disease-associated proteins (23% and 26%, respectively). We used k-means (k=6) to define metabolite/lipid clusters – these clusters showed enrichment of specific classes of lipids and metabolites (c). The protein clusters were enriched for specific genera (d). Protein clusters 2 and 3 had significant enrichment for GO-terms (e). TG, triglycerides; PE-NMe, mono-methyl phophospotidlyethanolamine; PE, phophospotidlyethanolamine; PG, phosphotidlyglyercol; SM, sphinomylin; PC, phosphotidylcholine; Cer[BS], ceramides containing beta-OH fatty acids and sphingosines; PD, periodontal disease; DM, diabetes.

To better explore the proteins and metabolites/lipids that composed the clusters, we preformed enrichment analysis (**Figure 5c-d**). We hypothesized that the protein clusters would contain proteins within related metabolic pathways; however, we found that most clusters were enriched in specific genera, while only a few clusters were enriched for specific pathways (GO biological processes, **Figure 5e**). This finding suggested that microbial populations, rather than specific pathways, had a stronger association with metabolite and lipid levels. Notably, protein clusters 2 and 3, which had the greater number of disease-associated features, were not enriched in bacterial proteins, but instead were enriched in human-derived proteins. These human-derived proteins were also associated with GO terms related to protein binding (cluster 2 and 3), enzyme regulator activity (cluster 2), actin binding (cluster 3), receptor binding (cluster 3), and lipid binding (cluster 3). Significant elevation of the actin-binding proteins in plaques from PD vs. non-PD patients is possibly due to bacterial invasion process and the loss of structural integrity at the tooth-endothelial interface ^40,41^. These proteins were also strongly correlated with abundance of many plasmenyl-PCs, PCs, and Cer[NS], suggesting these lipids might also be related to the host’s response to disease.

Many of the microbial-associated protein clusters in **Figure 5a** showed positive correlations with lipid/metabolite cluster 3 that was enriched in PE, PE-NMe, PG, and Cer[BS] lipids. As noted earlier, the PGs are likely derived from bacterial populations, and these clustering results are consistent with that conclusion. Likewise, PE-NMe are intermediates in the synthesis of PC in bacteria ^42^ and as such, would be expected to have association with bacterial proteins.

Further, protein cluster 1 was enriched in proteins derived from known oral pathogens: *Prevotella, Fusobacterium, Tannerella*, and *Selenomonas* genera ^9^. This large cluster exhibited positive correlations with many metabolites and lipids, *e.g.*, 5-aminovaleric acid, L-homoserine, hydrocinnamic acid, Cer[BS] containing odd-chain fatty-acyl chains, and Plasmenyl-PEs. In particular, 5-aminovaleric acid, a bacterial-derived metabolite generated during degradation of lysine, was positively correlated with proteins from *Selenomonas* and *Fusobacterium* – allowing us to hypothesize that these microbes might produce this metabolite. In general, the association between elevated amino acids and oral pathogens could explain why amino acids are biomarker candidates for PD ^38,39^.

Protein Cluster 6, which featured early biofilm colonizers like *Actinomyces, Corynebacterium*, and *Streptococcus*, displayed higher correlations with PE-NMe, malic acid, and 2-isopropylmalic acid. 2-isopropylmalic acid was only correlated with *Corynebacterium*, which reassuringly was the only genus with detectable protein levels of the necessary synthetic enzyme 2-isopropylmalic acid synthase (EC 2.3.3.13).

PE-NMe have been estimated to occur in only about 10-15% of bacteria ^32,43^, and in our analysis they were strongly correlated to *Lautropia*, a genus not previously described as synthesizing PE-NMe (**Figure 6a-b**). To validate this potentially novel finding, we compared lipid levels from two *in vitro* grown strains of *Lautropia mirabilis* to *Actinomyces odontolyticus. Actinomyces* had demonstrated low correlation to PE-NMe in our analysis, and thus was selected as a control. We confirmed PE-NMe synthesis in *Lautropia* and in fact found that PE-NMe were one of the more abundant lipids in these bacteria (**Figure 6c**).

**Figure 6.**
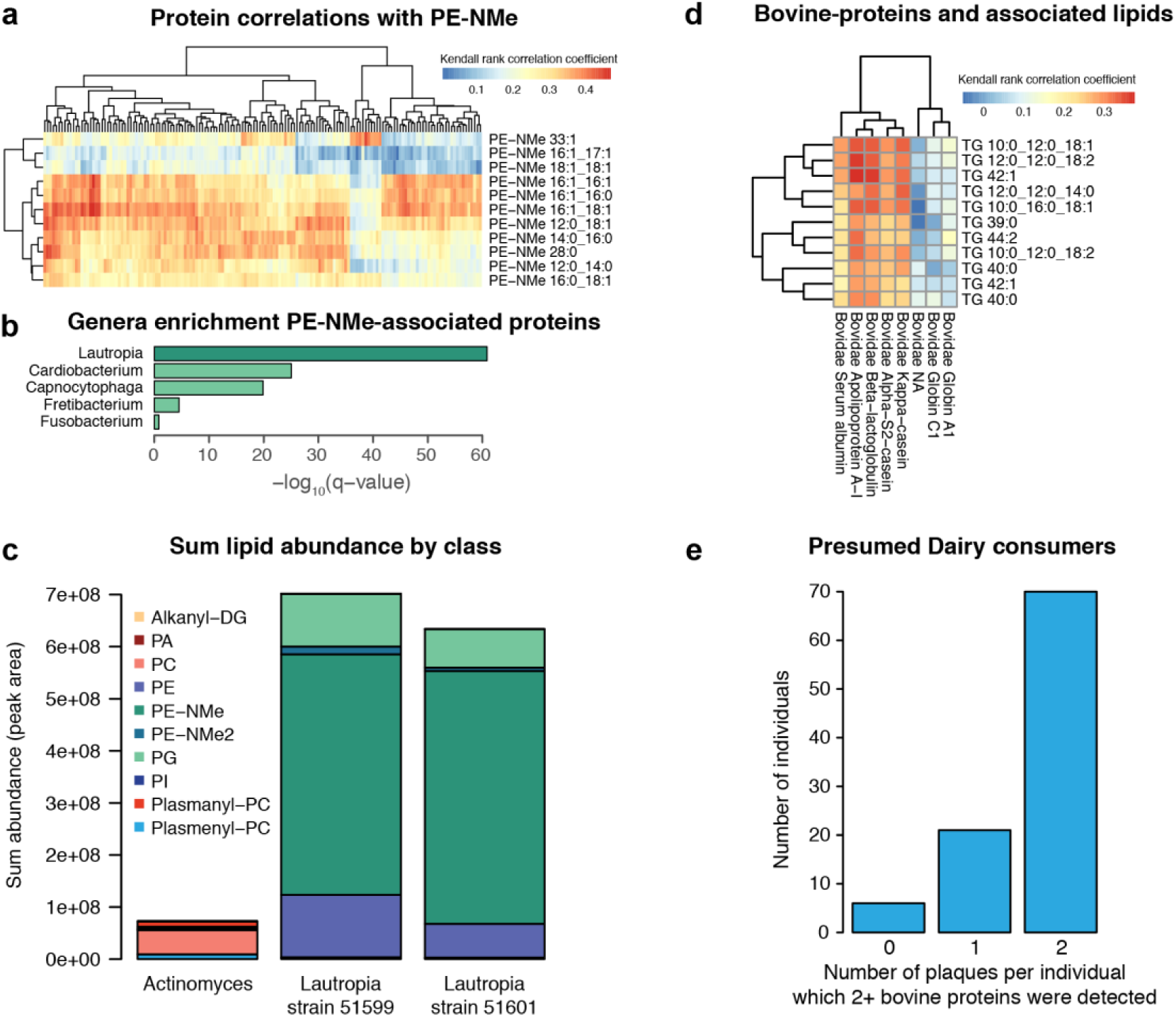
Lipid-protein associations facilitate observations about food consumption and microbial lipid synthesis pathways. Following from **Figure 5**, Kendall rank-based correlation was used to filter associations between metabolites, lipid, or proteins. PE-NMe were strongly associated with many bacterial proteins in protein cluster 6 of **Figure 5** (a). The PE-NMe associated proteins were highly enriched for *Lautropia* genera (b). Lipidomics profiles from single cultures of *Lautropia mirabilis* strains show PE-NMe are the dominate lipid class in these species (c). One small cluster show strong association between medium-chain length TG and bovine proteins (taxonomic family *Bovidae*) (d). Greater than 70% of individuals had dairy-associated proteins (2+ bovine proteins observed) in 2/2 plaque samples (e). TG, triglycerides; PE-NMe, mono- methyl phophospotidlyethanolamine; PE-NMe2, di-methyl phophospotidlyethanolamine; PE, phophospotidlyethanolamine; PG, phosphotidlyglyercol; PA, phosphotidic acid; DG, diacylglycerides; PI, phosphotidylinostitol, PC, phosphotidylcholine.

Finally, beyond host and microbial lipid associations, we found evidence of diet-associated features. In Protein Cluster 6 we found a lipid-protein association indicative of cow’s milk. One protein (Apolipoprotein A-I, of taxonomic family *Bovidae*) was positively correlated with several triglycerides containing medium-chain fatty-acyl tails (TG 12:0_12:0_14:0, TG 10:0_12:0_18:1, TG 10:0_16:0_18:1, TG 10:0_12:0_18:2, TG 12:0_12:0_18:2). The next most closely correlated proteins with these TGs were also assigned to taxonomic family *Bovidae* (alpha-S2-casein, kappa-casein, and beta-lactoglobulin). Together with APO-A1, these proteins constitute some of the most abundant proteins in cow’s milk ^44^. We annotated eight proteins to the taxonomic family *Bovidae*, and five of the eight showed strong correlations to medium-chain containing TGs (**Figure 6d**). As TGs are also highly abundant in cow’s milk ^45^, we concluded that this association was likely due to protein-lipid associations indicative of dairy consumption. Over 70% percent of the individuals had detectable levels of two or more of these bovine proteins in both of their plaque samples (**Figure 6e**).

In summary, the integrative MS-based multi-omics component of this analysis revealed findings beyond those typically seen with sequencing approaches, and facilitated discovery of host-disease, microbial-lipid, and diet-induced associations in dental plaques, associations which are critical to furthering our understanding of the human microbiome.

## Discussion

This study provides a comprehensive and comparative analysis of the microbiome, proteome, lipidome, and metabolome of dental plaque samples from individuals with PD and Pre-DM/DM. We detected on average 5,277 features per plaque from 97 individuals representing three disease groups and a control group, with over 7% of the detected features having disease associations with either gingival or metabolic health. We demonstrated that microbial dysbiosis occurred with PD, and that these changes were distinct from the microbial dysbiosis that resulted from Pre-DM/DM. Specifically, PD patient samples contained reduced levels of *Streptococci* relative to controls, and the Pre-DM/DM patient samples had increased abundance of *Fusobacterium* and *Tannerella*. We compared microbial population estimations obtained via 16S sequencing and proteomics. In general, we observed good agreement between the methods; however, some genera had higher estimations by 16S sequencing approaches (*Prevotella, Selenomonas*, and *Veillonella*), while other genera had higher estimations by proteomics (*Actinomyces* and *Corynebacterium*). Lastly, we used our MS data to correlate plaque proteins with metabolites and lipids. We revealed many microbial-molecule associations and importantly, discovered host-specific disease features, like actin filament-related proteins, which were highly correlated with PC and plasmenyl-PCs and strongly associated with PD. In sum, this study provides a data-rich multilayered analysis of the complex disease-associated ecosystem.

One of our findings was the observation of unique dysbiosis occurring with PD and Pre-DM/DM. Though PD is often a comorbidity of DM ^6,29^, we established that the supragingival microbial populations were distinct between PD and DM. In patients with PD, we observed lower relative abundance of *Streptococci*, a result consistent with the previous reports on reduced abundance of specific *Streptococcus spp*. in PD ^27,28^. This loss of *Streptococci* has been suggested to contribute to disease progression by freeing space for more pathogenic bacteria to thrive ^27^. In this study we did not specifically sample plaques from diseased tooth sites, thus it is possible this loss of more neutral bacteria is wide-spread in mouths of PD patients. In patients with Pre-DM/DM we observed elevated levels of periodontopathogenic pathogens, *Fusobacterium* and *Tannerella*, which also have been detected in other DM and non-DM obese populations ^29,46^. Specifically, Aemaimanam et. al. ^29^ reported on higher populations of these pathogens at healthy tooth sites in DM as compared to those in patients with PD alone, and that those microbial populations were correlated with HbA1c values, commonly used to monitor long-term glycemic control. This suggests that establishment of pathogenic bacteria at healthy sites correlated with systemic glucose load and might expedite progression towards PD.

Further, by utilizing MS-based multi-omics technologies, we discovered host-associated disease signatures. Plaques from PD patients contained significantly more host-derived proteins, which were enriched in actin-filament related proteins and likely have a mechanistic link to microbial invasion ^40,41^. With the goal to understand how microbiota contribute to disease, approaches like metaproteomics and metatranscriptomics, which enable detection of host response, will be critical for understanding these pathogenic interactions. We argue that MS-based ‘omics technologies could further strengthen our understanding of the host-microbe interactions, not only in being able to monitor potential protein response to microbial invasion, but also in monitoring lipid changes in these host-microbe environments. Lipids are important biomolecules related to defense and invasion as they reside at the interface between cells and serve as structural or signaling molecules ^43^. In support of this idea, we found that the same host-derived proteins associated with PD were also strongly correlated with lipids classified as PCs or plasmenyl-PCs. Further mechanistic studies will hopefully reveal how these molecules change during invasion and provide candidates for therapeutic intervention.

Beyond host-microbe interactions, lipidomics offers other novel insights about the oral microenvironment. A few recent studies have investigated saliva lipid profiles in chronic periodontitis ^20,47^, however, quantified only a small number of lipids. The current study offers a far more comprehensive profiling of the oral biofilm lipidome and provides an important first step in linking these lipids to taxonomic branches, diet, and oral health. The fact that this multi-omic approach revealed a true, and yet described, lipid pathway in *Laurtropia*, as well as a diet-related cluster suggestive of dairy consumption, showcases the discovery potential of this methodology. With the goal to better understand microbial microenvironments, a proteomic-lipidomic approach, like the one presented here, would offer significant biological insight, and we expect that future improvements to the methodology (i.e., improved time or space resolution) could be hugely beneficial to our understanding of this complex system.

## Methods

### Materials

Unless otherwise stated, materials were obtained from Sigma-Aldrich. Organic solvents and water used for extraction and MS-analysis were of MS-grade quality.

### Recruiting patients and sample collection

This study was approved by the IRB of Marshfield Clinic Research Institute under the IRB Protocol # SHU10115. PD and pre-DM/DM patients and healthy controls were recruited from the Marshfield Dental Clinic, Marshfield, WI based on their prior medical and dental records; in total 99 participants consented to the study; 2 patients were later excluded due to having a type I diabetes diagnosis. Participants were classified as Pre-DM/DM if they had been previously diagnosed with diabetes in their medical record or if they met the following criteria: fasting blood glucose 100 mg/dl or greater, HbA1C 5.7% or greater, and glucose tolerance test 140 mg/dl or greater. Patients were classified as having PD if they had undergone a periodontal exam and were classified as having moderate or severe periodontitis. Moderate periodontitis was classified as having either interproximal attachment loss >= 4 mm (2 or more teeth) or interproximal probe depth >= 5 mm. Severe periodontitis was classified as having both interproximal attachment loss >= 6 mm (2 or more teeth) and interproximal probe depth >= 5 mm. For every participant, supragingival plaque samples were collected from three locations, lower and upper molars and lower anterior lingual surfaces were targeted for collection, sample location was recorded by tooth numbers and surface (palatal vs. lingual), and samples were frozen in a dry ice-isopropanol bath within five minutes of collection. Samples were maintained at less than −20°C prior to analysis.

### 16S rDNA sequencing

The V4 region of the 16S rRNA gene sequencing was performed by following the protocol published in ^48^. The Illumina pair-end reads of partial 16S rRNA sequences were used as input for the QIIME analyses ^48^; the analysis was performed in the follow steps. 1) All the pair-end reads were assembled in one fastq file with samples independently tagged with their samples names, basic quality control steps were applied to make sure the quality of the fastq file, the parameters used are default of QIIME pipeline from http://qiime.org/tutorials/index.html. 2) OTU picking step was performed using the pick_open_reference_otus.py protocol by searching reads against the Greengenes database with similarity set to 99% ^49,50^. 3) Taxonomy assignment was performed using the ‘uclust’ method ^51^ and a 0.7 confidence cut-off with Greengenes taxonomy assignment ^35^. 4) Chimeric sequences were detected using the identify_chimeric_seqs.py function with the ‘usearch61’ method ^51,52^; these chimeric sequences were removed from the OTU table. The OTU results were exported as a ‘biom’ file and imported into R for further analysis.

### Bacterial culture preparation for lipidomics

Two ATCC strains of *Lautropia mirabilis* (ATCC strain #s 51599 and 51601) and a clinical isolate of *Actinomyces odontolyticus* were grown 5 ml tubes of Muller-Hinton Broth (MHB) at 37°C in static culture for 48 hours. After 48 hours, bacterial cells were centrifuged, supernatant discarded and fresh 5 ml of the MHB were added in to tubes, vortexed and incubated for additional 48 hours before collecting the cells pellet. All three cultures were grown in triplicates, and cell pellets were stored at −80°C prior to lipid extractions.

### Sample extraction for MS analysis

Samples were analyzed in batches of 25 with a cultured mixed-microbial quality-control sample in each batch. Samples were kept on dry ice prior to extraction. To each sample, we added 500 µL of ice cold extraction buffer (2:2:1 Methanol:Acetonitrile:Water). Samples were probe sonicated for 10s over ice and then centrifuged for 5 min at 14,000xg at 4°C to pellet protein and other debris. Supernatant was centrifuged again at 14,000xg for 5 min at 4°C to ensure no precipitate. The extract was divided for LC-MS-based lipidomics and GC-MS based metabolomics analyzes and dried by vacuum centrifugation. The precipitated protein was used for proteome analysis.

### GC-metabolomics

Dried extracts were resuspended in methoxyamine HCl (20 mg/mL in pyridine) and incubated at room temperature for 90 min. Samples were further derivatized at with MSTFA (Restek) for 30 min at 37°C. Samples were analyzed on a Q Exactive GC-Orbitrap mass spectrometer using a TraceGOLD TG-5SilMS GC column ^53,54^. Samples were injected using a 1:10 split at 275°C and ionized using electron impact (EI). The GC gradient ranged from 50- 320°C, linear over a 25 min gradient, then a 4.4 min hold at 320°C. Orbitrap MS-acquisitions were collected in full scan mode 50-650 m/z at a resolution of 30,000 (@ 200 m/z). Raw files were analyzed using an in-house tool for deconvolution of spectra, quantitation, and identification against in-house and NIST 2014 libraries ^55,56^.

### LC-lipidomics

Dried extracts were resuspended in 65:30:5 Isopropanol:Acetonitrile:Water. Samples were injected onto a Water’s Acquity UPLC CSH C18 Column (2.1mm x 100 mm) with a 5mm VanGuard Pre Column. Mobile Phase A: 70:30 ACN:H_2_O 10 mM NH4Ac 0.025% acetic acid. Mobile Phase B: 90:10 IPA:ACN 10 mM NH4Ac 0.025% acetic acid. The samples were run on a 30 min gradient. Mass spectra were acquired using a Thermo Focus Q-Exactive with polarity switching and top-2 data dependent ms2 scans. Raw files were analyzed with the Thermo Compound Discoverer™2.0 application with peak detection, retention time alignment, and gap filling; features were identified using LipiDex ^57^.

### LC-proteomics

Precipitated protein was solubilized in 8M urea prepared in 50 mM Tris, pH 8.0. Proteins were then reduced and alkylated with TCEP (10 mM final) and 2-chloroacetamide (40 mM final) for 15 min at room temperature, with shaking. Samples were diluted with 50 mM Tris, pH 8 to a final 4M urea concentration, then proteins were digested overnight with endoproteinase Lys-C (1:100 enzyme:protein, Wako Pure Chemical Industries). Samples were desalted with C18 SepPak columns (Waters); peptides were then dried down and resuspended in 0.2 % formic acid. Peptide concentration was estimated using a peptide colorimetric assay (Pierce), and 1 µg of peptides were analyzed by LC-MS/MS using a nano-LC column coupled to a Thermo Orbitrap Elite ^58^. Raw files were searched against a concatenated database containing peptides form the Human Oral Microbiome Database ^59^ and peptides in the human Uniprot database using a two search strategy ^60^. First, the initial search was completed individually on each sample using COMPASS ^61^, then we combined all first search identification matches to create a reduced fasta database for a second search using MaxQuant ^62^. Identified peptide sequences were queried against NCBI’s NR database (protein blast) ^63^ and resulting hits were filtered using MEGAN6 ^64^ to assign lowest common ancestor to each peptide, which we then assembled into functional and taxonomy assignment at the protein groups level generated by the MaxQuant algorithm.

### Data Analysis

Data were analyzed using the R statistical and graphing environment ^65^. Normalization for batch effects were done with ComBat ^23^. For statistical analysis, we modeled the effect of diabetes and periodontal disease on the abundance of each molecule with generalized additive mixed-effect models (GAMLSS ^66^). We used models with fixed effects for diabetes, periodontal disease, interaction between diabetes and periodontal disease, and confounding factors of age, gender, and tobacco-use status. To account for replicate sampling from individuals (in MS-acquired data), we included participant identifier as a random effect in the models. Due to the differences in analysis paradigms, we chose different data distributions to best fit the data: zero adjusted Gamma distribution (16S data and proteomics), log normal distribution (lipidomics), and bimodal log normal distribution (metabolomics, due to imputed values of low-level features We evaluated significance of Pre-DM/DM and PD on our models with log-likelihood ratio testing and Benjamini-Hochberg false discovery rate correction. For analysis of microbial diversity, we used R package vegan ^67^, and for plotting heatmaps we used pheatmap ^68^.

## Acknowledgments

We thank DeeAnn Polacek and Dixie Schroeder for their administrative support, and Evgenia Shishkova for assistance with editing the manuscript. We acknowledge the support of the NIH P41 GM108538. K.A.O., T.W.R., and A.E.M. were supported by NLM training grant 5T15LM007359. K.A.O. was supported through the Morgridge Institute for Research Postdoctoral Fellowship.

## Author Contributions

A.E.M., T.W.R., S.S., A.A., M.S.W., J.J.C. designed experiment. T.W.R., K.A.O., S.S., and Z.Y. performed data acquisition and analysis. All authors contributed to writing and editing manuscript.

